# Adopting Literature-based Discovery on Rehabilitation Therapy Repositioning for Stroke

**DOI:** 10.1101/422154

**Authors:** Guilin Meng, Yong Huang, Qi Yu, Ying Ding, David Wild, Yanxin Zhao, Xueyuan Liu, Min Song

## Abstract

Stroke is a common disabling disease severely affecting the daily life of the patients. There is evidence that rehabilitation therapy can improve the movement function. However, there are no clear guidelines that identify specific, effective rehabilitation therapy schemes, and the development of new rehabilitation techniques has been fairly slow. One informatics translational approach, called ABC model in Literature-based Discovery, was used to mine an existing rehabilitation candidate which is most likely to be repositioned for stroke. As in the classic ABC model originated from Don Swanson, we built the internal links of stroke (A), assessment scales (B), rehabilitation therapies (C) in PubMed relating to upper limb function measurements for stroke patients. In the first step, with E-utility we retrieved both stroke related assessment scales and rehabilitation therapies records, and complied two datasets called Stroke_Scales and Stroke_Therapies, respectively. In the next step, we crawled all rehabilitation therapies co-occurred with the Stroke_Theapies, named as All_Therapies. Therapies that were already included in Stroke_Therapies were deleted from All_Therapies, so that the remaining therapies were the potential rehabilitation therapies, which could be repositioned for stroke after subsequent filtration by manual check. We identified the top ranked repositioning rehabilitation therapy following by subsequent clinical validation. Hand-arm bimanual intensive training (HABIT) ranked the first in our repositioning rehabilitation therapies list, with the most interaction links with Stroke_Scales. HABIT showed a significant improvement in clinical scores on assessment scales of Fugl-Meyer Assessment and Action Research Arm Test in the clinical validation on upper limb function for acute stroke patients. Based on the ABC model and clinical validation of the results, we put forward that HABIT as a promising rehabilitation therapy for stroke, which shows that the ABC model is an effective text mining approach for rehabilitation therapy repositioning. The results seem to be promoted in clinical knowledge discovery.

**Author Summary:** In the present study, we proposed a text mining approach to mining terms related to disease, rehabilitation therapy, and assessment scale from literature, with a subsequent ABC inference analysis to identify relationships of these terms across publications. The clinical validation demonstrated that our approach can be used to identify potential repositioning rehabilitation therapy strategies for stroke. Specifically, we identified a promising rehabilitation method called HABIT previously used in pediatric congenital hemiplegia. A subsequent clinical trial confirmed this as a highly promising rehabilitation therapy for stroke.

## Introduction

### Stroke and rehabilitation

Stroke is a common disabling health-care problem, which is attributed to be the second-leading cause of mortality and disability worldwide [1,2]. For instance, in UK stroke is the largest single cause of disability with an annual cost to society of approximately £9 billion; in the United States, nearly 0.8 million people have stroke annually and the estimated direct and indirect costs of stroke is $95 billion in 2015, expected to rise to 185 billion in 2030 [3]. The symptoms of acute stroke include physical impairments and cognitive dysfunction, and physical impairments of the affected limbs range from movement restriction, sensory loss, muscle activation abnormalities, etc.[4]. About 50% acute stroke survivors suffered from dysfunction of the upper limbs in their chronic phase [5], severely impacting the daily life and the therapeutic effect of rehabilitation therapy, which reduces the quality of life after stroke [6,7].

Rehabilitation therapies offer a chance for an individual to recover and adapt to situation following acute stroke. There has been a large amount of research into methods of rehabilitation management, including task-oriented training [8], impaired limb forced training [9], movement science-based therapy, robotic-assisted movement, virtual reality (VR) training [10], functional electrical stimulation [11], and skill acquisition training paired with impairment mitigation and motivational enhancement and etc.

Currently, no high-quality evidence can be found for any rehabilitation management that is currently used as part of routine guideline practice, and evidence is not sufficient enough to evaluate the relative effectiveness of existing rehabilitation strategies in large clinical trials [12]. Furthermore, the stagnant development of new competitive rehabilitation strategies impedes rehabilitation therapy repositioning from possible unfolding data sources, such as large amount of literature in PubMed.

### Rehabilitation therapy repositioning based on Don Swanson’s ABC model

Rehabilitation therapy repositioning is the application of already approved therapy to new diseases, which is derived from Data-driven Method (DDM). DDM is based on analyzing data about a system, in particular finding connections between the system state variables (input, internal and output variables) without explicit knowledge of the physical behavior of the system [13]. Scientific literature is a special kind of data, usually semi-structured or unstructured, which comprises scholarly publications that report original empirical and theoretical work in the natural and social sciences, and within an academic field. Literature mining is a specialized data mining method that is used to extract information (facts or data) from scientific literature [14]. Applying DDM to literature mining can generate rehabilitation therapies, which can be repositioned for stroke by systematically scrutinizing a vast amount of abstracts, or full text versions of scientific publications.

The principal advantages of rehabilitation therapy repositioning over new therapy development are that approved therapy has already been tested for safety, and repositioning can eliminate the time and cost of developing new therapy. For instance, the use of VR as a rehabilitation intervention was first applied to basic motor disability [15]. Research in the area of VR based rehabilitation gained growing recognition of the potential value of VR for other diseases with motor disorders, for example, Parkinson’s disease. VR is now proposed as a new rehabilitation tool that potentially optimizes motor learning in a safe environment and replicates real situations to help improve functional activities in daily life such as gait, balance, and quality of life [16].

In order to identify new repurposed rehabilitation therapies for stroke, we developed a relation extraction method using ABC model proposed by Don Swanson [17]. ABC model demonstrated that new knowledge could be discovered from sets of disjointed scientific articles (Fig 1). In the model shown in Fig 1, one set of articles (AB) reports an interesting association between variables A and B, while another set of articles (BC) reports a relationship between B and C, but nothing at all has been published concerning a possible link between A and C, even though such a link if validated would be of scientific interests. The open ABC model is to start with theme “A” in MEDLINE or PubMed that collects scientific questions (such as literatures that discuss stroke); the word / phrase “B” is then listed (in the title or abstract appearing in A) and a separate literature search is used for each “B” term (or filtered subset); then the words and phrases “C” appearing in the code of the B are compiled (or filtered); finally, by some criteria, the C terms are ranked such that a high ranked C term is said to represent the most promising hypothesis. Depending on the system, B and C may represent other features extracted from medical topic titles or concepts. For example, the term C may be the name of a therapy strategy that has not been tested for stroke for A (stroke), but C (rehabilitation therapy) has been demonstrated in other situations (e.g., in other forms of physical injury model or experimental animal model) with curative effects, suggesting that C may be explored as a new therapy. For each disease, there should be corresponding therapies including rehabilitation therapies; meanwhile, for each rehabilitation therapy, there should be corresponding measurements, most of which are assessment scales focused on different aspects of rehabilitation effectiveness. So the identification of disease – rehabilitation therapy, rehabilitation therapy – assessment scale and disease – rehabilitation therapy relationships is key to identifying and curating new candidates for rehabilitation repositioning based on the ABC model.

**Fig 1.**
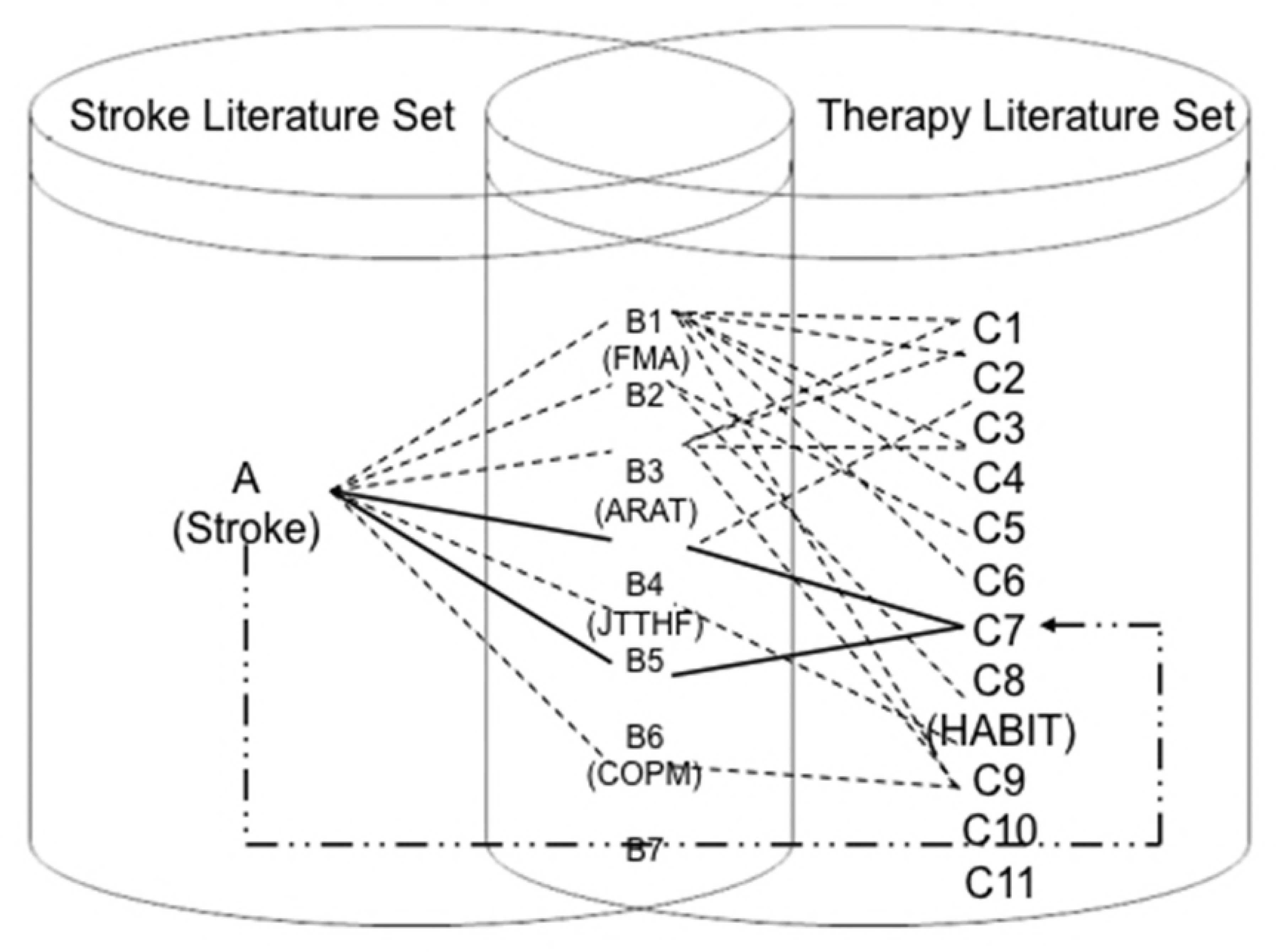
ABC model of Stroke-Assessment scales-Rehabilitation therapies

Since there are no databases that extensively curate the relationships among these three entities, unidentified relationships may be buried in literature [18,19]. Recently, literature mining has been applied to automatically generate various, plausible new hypotheses [20,21], which motivates to find new indications of existing rehabilitation therapies for stroke.

In this study, with the basis of ABC model, we aimed to detect indirect relationships that may facilitate the discovery of new candidates supporting the curating for rehabilitation repositioning. Specifically, the aim of this study is to find new rehabilitation therapy for stroke by extracting relationships from the literature and to provide clinical validation to identify the most promising potential therapy.

## Results

### Text mining based on ABC model

We retrieved 11,418 records from PubMed with stroke related keywords, and there were 10,992 records with abstract. From this dataset, 241,044 unique NPs were extracted, which included 81 unique scales and 215 unique rehabilitation therapies.

In the potential scale list, the common phrases such as “pre-test”, “post-test”, “outcome assessment” as well as other unrelated scales such as “body mass index” and “depression score” and “MMSE score” were deleted after manual check from the stroke related scale dataset, Stroke_Scales, which ended up 28 scales (Fig 2).

**Fig 2.**
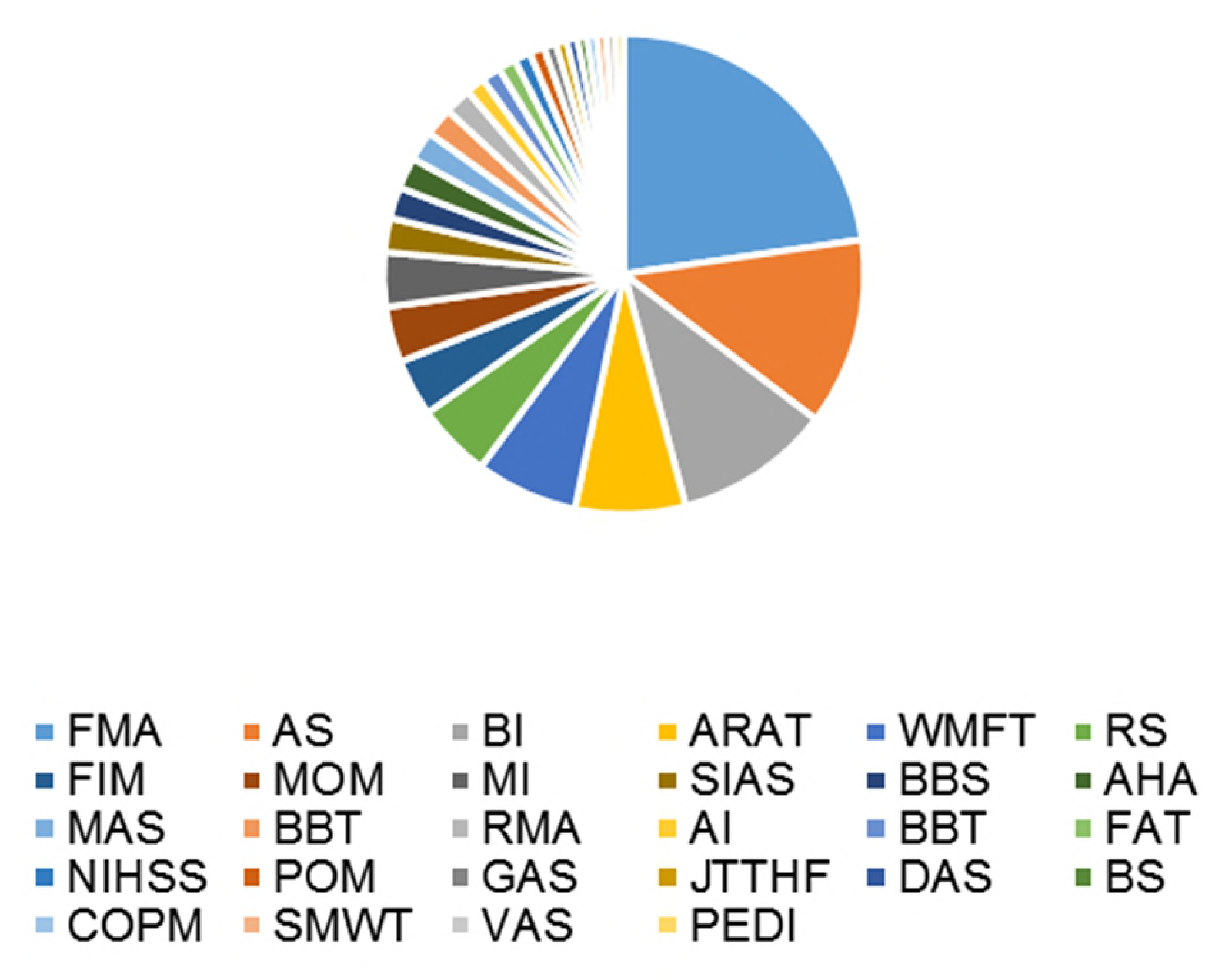
Stroke_Scales dataset items with frequency.

FMA (fugl - meyer assessment, AS (ashworth scale), BI (barthel index), ARAT (action research arm test), WMFT (wolf motor function test), RS (rankin scale), FIM (functional independence measure), MOM (main outcome measure), MI (motricity index), SIAS (stroke impact scale), BBS (berg balance scale), AHA (Assisting Hand Assessment), MAS (motor assessment scale), RMA (rivermead motor assessment), AI (arm index), BBT (box and block test), FAT (frenchay arm test), NIHSS (National Institute of Health stroke scale), POM (primary outcome measure), GAS (Goal Attainment Scale), JTTHF (Jebsen-Taylor Test of Hand Function), DAS (disability assessment scale), BS(brunnstrom scale), COPM (canadian occupational performance measure), SMWT (Six-Minute Walk Test, VAS (visual analogue scale), PEDI (Pediatric Evaluation of Disability Inventory).

The FMA had the largest share of 22.7%, followed by AS, BI, ARAT, WMFT, RS, FIM, and MOM, whose total share was 72.8%, which means the most widely used assessment scales in stroke related studies.

Accordingly, in the potential therapy list, the following common words were deleted: “clinical practice”, “conventional therapy”, “medical therapy”, “physical therapy”, “specific training”, and “combined therapy” and so on; in addition, the following drug therapy of “antiplatelet therapy”, “anticoagulant therapy”, “antihypertensive therapy”, and “antithrombotic therapy” were deleted. Thus, we got the stroke related rehabilitation therapy dataset, Stroke_Therapies, which compromising 47 rehabilitation therapies (Table 1).

**Table 1.**
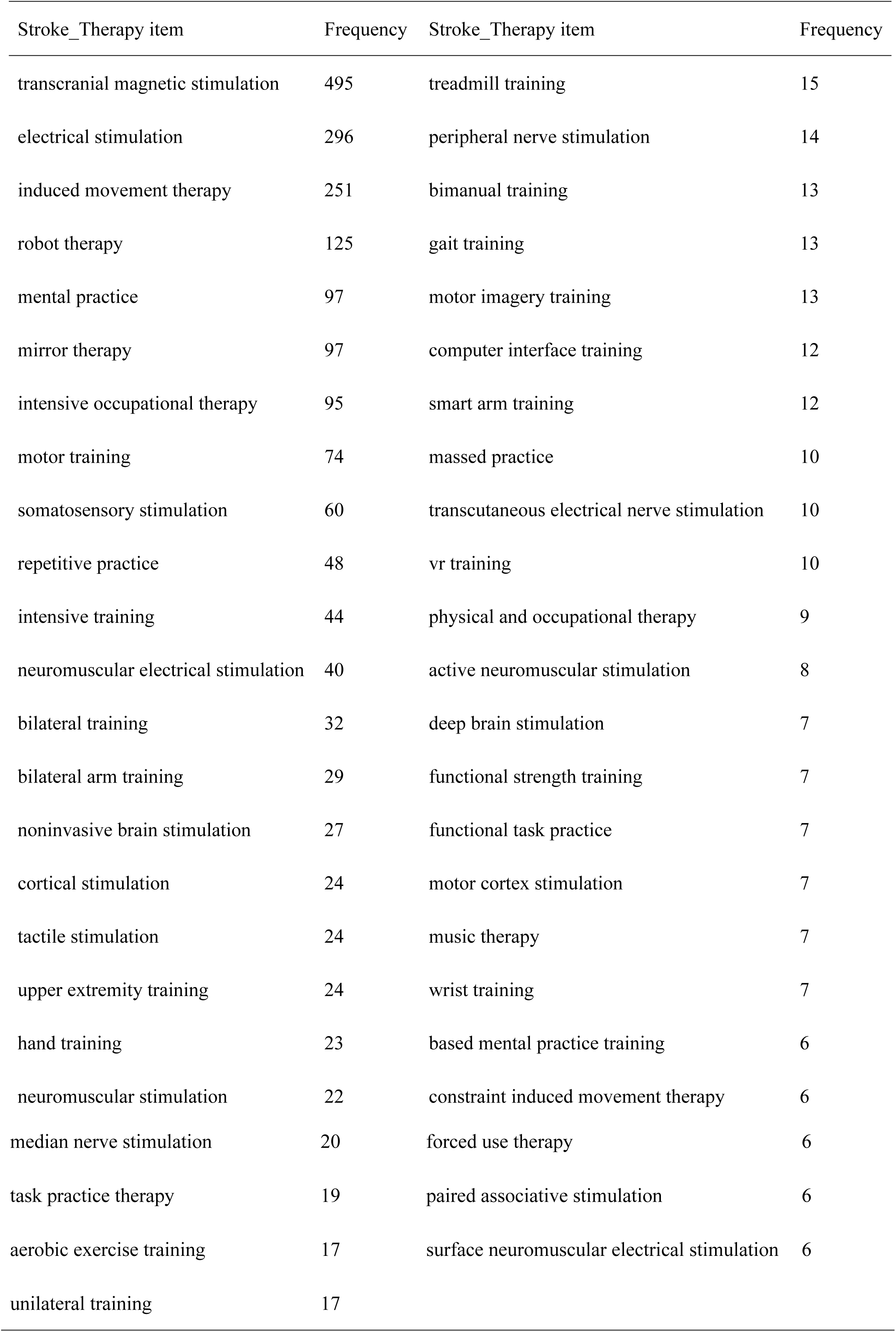
Stroke_Therapies dataset items matching with frequency

With the extracted Stroke_Scales, we retrieved 60,307 records in which 60,202 records with abstract in PubMed. From these records, we extracted the rehabilitation therapies (All_Therapies dataset), and removed those that have been applied for stroke, which is listed in Table 1 (Stroke_Therapies dataset) and got the potential repositioning rehabilitation therapies for stroke in Table 2. The interactions of Stroke_Scales and All_Therapies dataset were shown in Fig 2.

**Table 2.** Potential repositioning rehabilitation therapies

| Unknown rehabilitation therapies | Frequency | Unknown rehabilitation therapies | Frequency |
| --- | --- | --- | --- |
| Cognitive behavior therapy | 146 | hand arm bimanual intensive training | 7 |
| massage therapy | 30 | intensive bimanual training | 2 |
| homeopathic treatment | 16 | home based intensive bimanual training | 2 |
| acupuncture treatments | 12 |  |  |

We used these potential repositioning rehabilitation therapies with “stroke” in PubMed search to exclude the records that contain the associations of those rehabilitation therapies with stroke. We found that except for “hand arm bimanual intensive training” and “home based intensive bimanual training”, the rest of NPs co-occur with stroke. Thus, those two previously unknown rehabilitations, “hand arm bimanual intensive training” and “home based intensive bimanual training”, worth further investigation. With the consideration of the practical situation where rehabilitation is started in the acute phase of stroke patients, it is apparent that our target population were acute stroke patients treated in hospital but not at home, so that “home based intensive bimanual training” was not applied in this study, we took the “hand arm bimanual intensive training” as the first choice for clinical validation.

In the second quadrant of Fig 3, Stroke_Scales were densely distributed and were interacted with All_Therapies dataset from other three quadrants. Among them, the potential repositioning rehabilitation therapies were marked in red and rose, while the existing Stroke_Therapies dataset items were indicated in other colors. Among the potential repositioning rehabilitation therapies, “hand arm bimanual intensive training” shared the most interactions with Stroke_Scales, including Jebsen-Taylor Test of Hand Function (JTTHF), Canadian Occupational, Performance Measure (COPM), Assisting Hand Assessment (AHA), Pediatric Evaluation of Disability Inventory (PEDI), Box and Blocks Test (BBT), Six-Minute Walk Test (SMWT), Goal Attainment Scale (GAS), none of which was the commonly used stroke related assessment scales according to targeting populations and disciplines difference.

**Fig 3.**
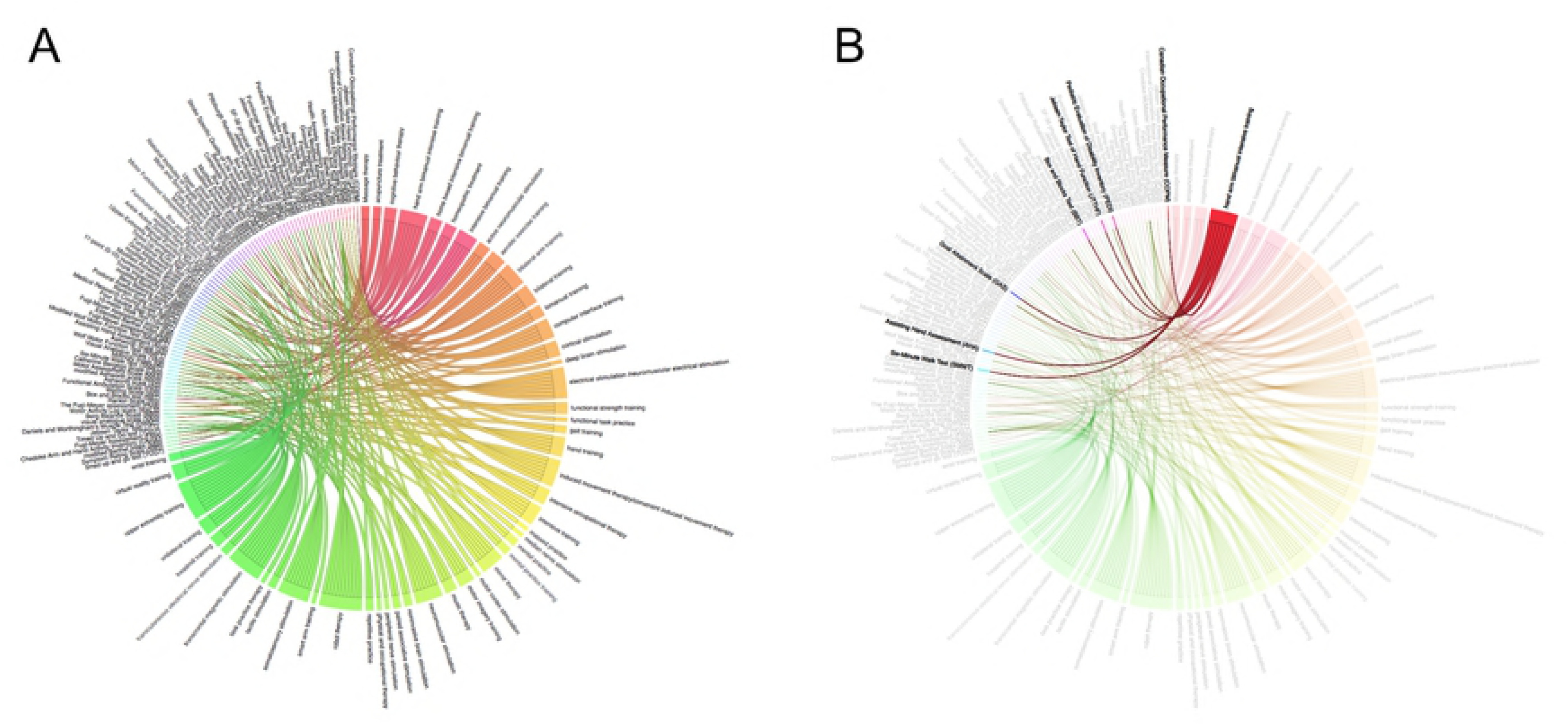
A. Interactions of assessment scales and all therapies; B. Interaction of HABIT and assessment scales.

### Hand–arm bimanual intensive training (HABIT)

HABIT is a bimanual intervention addressing the specific upper extremity impairments in pediatric congenital hemiplegia, which is the most common physical disability in childhood [22,23], typically with impairments of spasticity, sensation, and reduced strength. HABIT has been reported to improve the pediatric patients bimanual hand coordination and the space control of actions. Furthermore, HABIT is demonstrated to be the prioritized optimal approach to improving bimanual hand use and activity performance for children with hemiplegia [24], whose principles include motor learning (practice specificity, types of practice, feedback) [25], and principles of neuroplasticity (practice-induced brain changes arising from repetition, increasing movement complexity, motivation, and reward) [26,27], which are also the key contents of stroke rehabilitation functional goals.

We checked the frequency of different scales for stroke, and found that compared with JTTHF and COPM, Fugl-Meyer assessment (FMA) was the most commonly used measure for the upper limb in adult populations, almost 30% of the total frequencies (Table 1), which is also in accordance with a systematic literature about stroke rehabilitation studies [28]. Based on the fact that FMA was usually applied in combination with action research arm test (ARAT) in clinical studies, we decided to apply FMA combined with ARAT, not COPM or JTTHF in our clinical validation.

The final validation of the potential candidate of repositioning rehabilitation therapy was conducted in the clinical trial of stroke patients with upper limb impairment. The trial was registered with ChiClinicalTrials.gov, number CTR-INR-1701046 [29]. To our best knowledge, there have been no other clinical studies besides ours evaluating HABIT as a therapy for adult acute stroke patients. In general, patients were assessed by FMA and ARAT for motor function and extremity activity pre-and post-2 consecutive weeks of HABIT therapy. As shown in Table 3, after the rehabilitation therapy, a direct comparison of FMA and ARAT scores revealed the significant improvement, showing that HABIT improved the scores in both scales.

**Table 3.** FMA and ARAT scores pre-and post-HABIT rehabilitation therapy

| Scales | Pre-therapy | Post-therapy | *P* |
| --- | --- | --- | --- |
| FMA | 33.25±5.89 | 51.73±6.44 | <0.001 |
| ARAT | 30.31±6.07 | 34.47±6.22 | <0.001 |

## Discussion

### ABC model in rehabilitation therapy repositioning

Knowledge is sometimes segregated by syntactically impenetrable keyword barriers or undiscovered in an entirely different research corpus, so that clinicians in Neurology domain may not be able to keep completely up-to-date with applicable rehabilitation therapies for stroke patients. Analyzing the literature and data to semi-automatically generate a hypothesis about rehabilitation therapy repositioning might become the de facto approach to inform clinicians who are trying to master the exponentially rapid expansion of publications and datasets. Swanson [17] proposed the ABC model that can be applied to new hypothesis generation for rehabilitation therapy repositioning, where ABC model is pertinent to an association rule between a separate set of publications: if A is associated with B, and B is associated with C, then there is a potential relation between A and C. ABC model has played a number of roles in the direction of drug discovery and repositioning. Earliest applications of the ABC model derived from two major findings of fish oil treatment for Raynaud’s disease and magnesium treatment for migraines, both of which have been clinically confirmed [30]; in recent years, vigorous development of bioinformatics mining and omics study indirectly borrowed the mode l [31]. However, one biggest limitation of the current ABC model-based approaches is that without certain domain knowledge, it is not easy to identify significative AB and BC directly. In our study, the neurology clinicians (domain experts) were fully engaged, including raising questions, checking manually and confirming the final candidate on the basis of expertise, which eliminated the limitation

On the other hand, although the repositioning of rehabilitation therapy is not uncommon in clinical practice, this discovery is often based on the clinician’s personal experience or deduction from the medical community, without objective, systemic approach to data mining application. In our discovery case, stroke (A) is associated with the assessment scales (B) in stroke literature, and in the rehabilitation therapy literature, assessment scales (B) represent the effect of rehabilitation therapy (C), then, it is highly likely that the retrieved rehabilitation therapies that are unknown for stroke yet have a positive effect on stroke. It is the first application of ABC model to show the repositioning of rehabilitation therapy with positive validation. This model could be generalized as disease-assessment scale-rehabilitation therapy in future studies.

### HABIT helps upper extremity function recovery for stroke

In the clinical validation, the patient group who received HABIT demonstrated significant improvement indicating that HABIT has a positive impact on rehabilitation therapy for upper extremity impaired patients.

The principle of HABIT includes motor learning (task specificity, task type and feedback) and neuroplasticity, as well as brain transformation upon increasingly difficult therapy and incentive reward [26], training cortico-spinal system reconstruction, manifesting as function recovery after injury [32], which is the theoretical foundation of hypothesizing that bimanual intensive therapy could improve the upper extremity function after acute stroke.

In the present study, we proposed a text mining approach to mining terms related to disease, rehabilitation therapy, and assessment scale from literature, with a subsequent ABC inference analysis to identify relationships of these terms across publications. The clinical validation demonstrated that our approach can be used to identify potential repositioning rehabilitation therapy strategies for stroke. Specifically, we identified a promising rehabilitation method called HABIT previously used in pediatric congenital hemiplegia. A subsequent clinical trial confirmed this as a highly promising rehabilitation therapy for stroke. As a follow-up study, moreclinical trials should be involved for the long-term impact of HABIT on stroke patients, and optimal parameterization of the therapy. We also plan to refine the text mining and inference strategy and apply to other clinical applications amenable to the technique.

## Materials and methods

### Study procedure

In ABC model, there is an internal connection among disease, rehabilitation therapy, and assessment scale. For stroke, most assessment scales of upper limb impairment rehabilitation assess a person’s ability to manage daily activities that require the use of the upper limbs, whatever the therapy strategies involved; meanwhile, those rehabilitation therapies for functional improvement of upper limb share the same set of scales for assessment, regardless of whether they are applicable of stroke or not yet. So our plan was to identify undiscovered rehabilitation therapies for stroke through shared assessment scales (B), and ultimately clinically validate the most promising candidate.

### ABC model implementation

Our first focus was to find repositioning rehabilitation therapy candidates from an extensive collection of articles in PubMed. The flow chart was shown in Fig 4.

**Fig 4.**
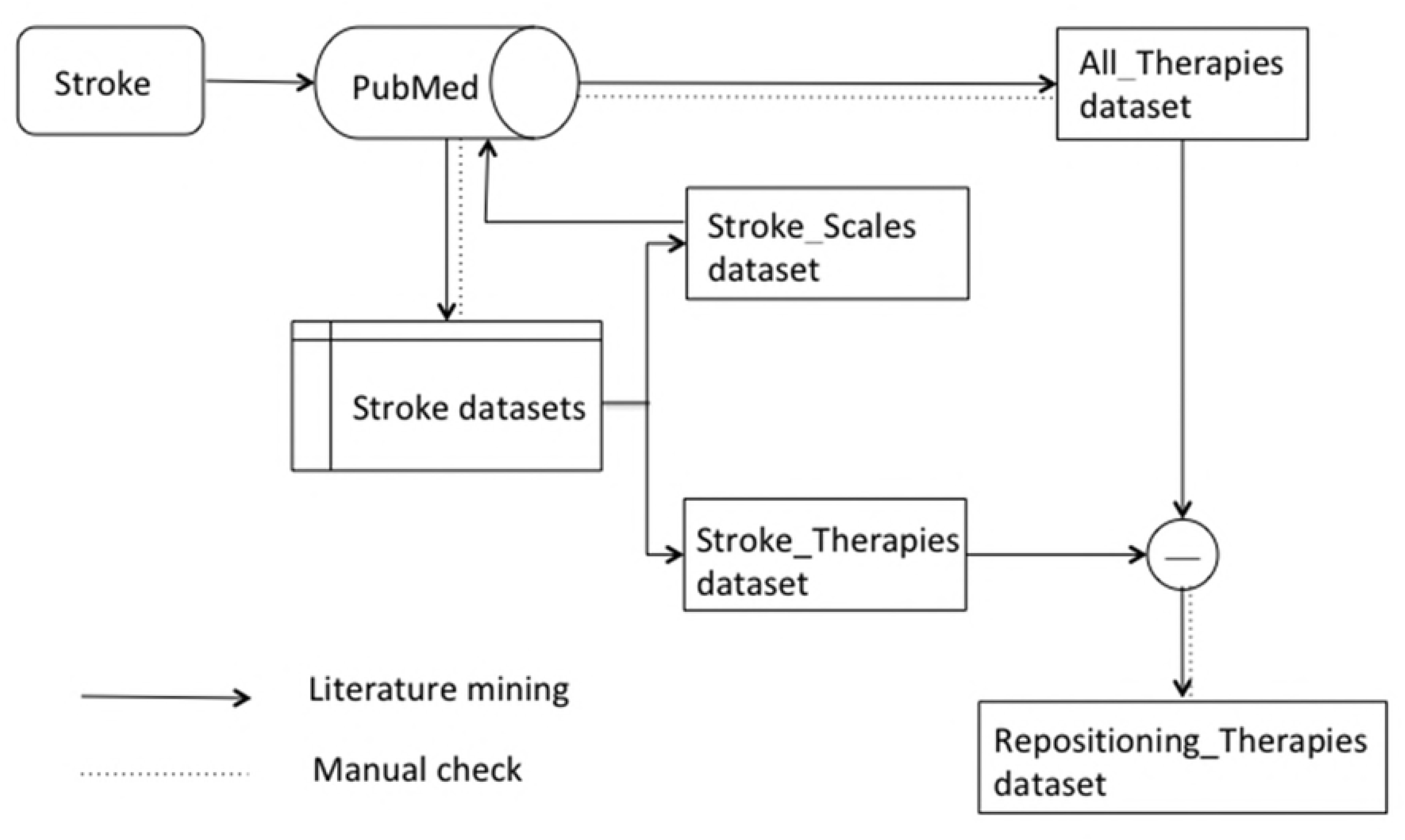
Flow chart from stroke to repurposed rehabilitation therapy

(The code of this article can be found in https://github.com/hyyc116/Stroke_findings/tree/master/DSTN.)

### Stroke related assessment scales and rehabilitation therapy datasets creation

To collect articles related to stroke with upper limb impairment, we searched PubMed with stroke related keywords: (“stroke” OR “cerebral infarction” OR “brain ischemia” OR “cerebral hemorrhagic” OR “subarachnoid hemorrhage”) and (“hand” OR “arm” OR “upper extremity” OR “upper limb”), with the proviso of humans.

The reason why we did not use Medical Subject Headings (MeSH) terms is that most specific assessment scales and therapies do not directly belong to MeSH terms. When we searched stroke related rehabilitation items, the MeSH terms we got were only the common words (see complementary material “MeSH terms”), which should be deleted as confounding factors.

E-utilities (https://www.ncbi.nlm.nih.gov/books/NBK25500/) were used to fetch all query related data in the PubMed. Scales and rehabilitation therapies are always noun phrases (NPs) in scientific articles. Thus, we applied a shallow chunk analyzer to extract NPs. The NPs ended with “test”, “scale”, “assessment”, “measure”, “score” or “index” with frequency more than 5 were kept to be a possible stroke assessment scale candidate. In addition, NPs ended with “training”, “therapy”, “treatment”, “treatments”, “practice”, “program”, “practise” or “simulation” with frequency more than 5 as well were kept to be a possible stroke therapy candidate.

Since relying only on the predefined lexicon to gather rehabilitation documents could lead to false negatives, manual inspection should be conducted to identify true positives as accurately as possible before stroke related scale dataset (Stroke_Scales) and rehabilitation therapy dataset (Stroke_Therapies) were established.

#### Entire rehabilitation therapy dataset creation

ABC model has been successful in explaining how two concepts are linked by an intermediate therapy discovery [17]. Specifically, with a direct stroke - assessment scales and therapies - assessment scales relationship, we crawled all therapies related NP in PubMed co-occurred with Stroke_Scales via E-Utility. The proceeding entire rehabilitation therapy dataset, named All_Therapies building was similar with Stroke_Scales building mentioned above, that the NPs ended with “training”, “therapy”, “treatment”, “treatments”, “practice”, “program”, “practice” or “simulation” with frequency more than 5 were kept to be a possible therapy candidate in which the assessment was the item in Stroke_Scales. Manual inspection was the same as above of Stroke_Therapies.

#### Potential repositioning rehabilitation therapies dataset

Therapies already included in Stroke_Therapies were deleted from All_Therapies, so that the remaining therapies were not stroke-applied therapies, which could be repurposed for stroke. Manual inspection was carried out again.

### Hypotheses validation

#### Validation of the retrieved Repositioning_Therapies dataset in PubMed

The articles related to potential repositioning rehabilitation therapies together with stroke related keywords were retrieved from PubMed to ensure that there are no articles that contain associations of therapies with stroke.

#### Further rehabilitation theory of potential candidate exploration

The knowledge discovery is full of uncertainty and complicated, in which algorithms and methods could be perfect in theory, while the precision, recall or some other metrics could be meaningless to some extends. Thus, what we aim at is to find potential candidates. Clinical value of the potential candidates then should be further comprehensively explored in the mechanism and principles with rehabilitation theory.

#### Validation in clinical trials

A clinical trial of adult acute stroke patients was carried out to test the rehabilitation effect by analyzing several sorts of data, aiming to clarify the potential advantage of rehabilitation therapy and to determine the optimal rehabilitation approach for stroke patients.

## Acknowledgments

This study was supported by the National Natural Science Foundation of China (81771133, 71573162) and also partly supported by the Bio-Synergy Research Project (NRF-2013M3A9C4078138) of the Ministry of Science, ICT and Future Planning through the National Research Foundation.

